# Prediction-Guided Design of a More Developable FGF21 Construct

**DOI:** 10.64898/2026.07.13.738140

**Authors:** Çağlar Bozkurt, Evangelia Nathanail, Aniruddh Goteti

## Abstract

For structural-biology and protein-production pipelines, the hardest part of a difficult protein is not the biology — it is obtaining a well-behaved sample for functional studies. Programs routinely stall at *construct design, expression, and purification*: deciding where to truncate, which tags to use, how to express, and how to purify so the protein survives concentration and handling. These decisions are still made largely by literature precedent and experimental experience, and they require trial-and-error before arriving at a functional construct for hard targets.

We present a prospective, single-pair wet-lab case study testing whether an integrated computational platform can improve these decisions. For human fibroblast growth factor 21 (FGF21) — a clinically important and stability-challenged metabolic hormone — we compared two expression constructs produced side by side under the same experimental workflow, using two different design strategies: one designed by a scientist from the literature (reproducing the published core-domain construct, PDB 6M6E), and one designed by the Orbion platform — an AI, prediction-guided protein-design system (orbion.life) — which additionally generated the expression and purification protocols (executed scientist-in-the-loop). The platform’s construct used an unconventional, longer C-terminal boundary not found in public sequence databases. Since the two constructs differ in more than one feature, we treat them as workflow-level designs throughout.

The scientist construct gave a higher *initial* yield (∼2.4 ×more protein recovered at affinity capture). The platform-designed construct, however, showed a more favourable downstream developability profile: it concentrated higher (1.4 vs 0.7 mg/mL) while remaining more monodisperse by dynamic light scattering (DLS). The scientist construct, in contrast, aggregated on concentration, so its initial-yield advantage did not survive: in the final concentrated sample the Orbion construct provided the more usable material for downstream studies. Computed for the mammalian host used, the platform had *prospectively* scored its own design higher (composite 68.7 vs 59.0 for the scientist-designed construct), and its predictions of yield, solubility, and disorder matched the wet-lab outcome.

This is a single, deliberately scoped case study, not a population-level benchmark; the two constructs differ in more than one feature, and biological activity was not assayed. Alongside the bottlenecks of this approach discussed here, used as a decision aid, prediction-guided construct and protocol design has the potential to remove costly iteration cycles of protein production campaigns.

## 1. Introduction

### The Upstream Bottleneck

Once a structure can be predicted confidently, working with a protein experimentally is often assumed to be straightforward. In practice, the rate-limiting step for difficult targets is earlier in the structural pipeline — obtaining well-behaved material at all: programs commonly stall at construct design, expression, and purification before the biology is tested [14, 15]. Constructs are screened largely by trial-and-error [13], with a substantial fraction failing to express. For many targets the limiting failure lies not in the biology but in the basic behaviour of the protein itself — poor stability, aggregation, low solubility, or loss on concentration.

These decisions — where to truncate, which tags, fusions, and signal peptides to use, which host to choose, and how to purify so the protein survives concentration — are made primarily from literature precedent and prior experience, which is least informative for novel targets with no prior structure or documented expression conditions. Reported success rates are correspondingly low: for membrane proteins the per-experiment failure rate exceeds 90% [16], and across large structural-genomics pipelines only a minority of cloned targets have historically yielded purified, well-behaved protein [14], though high-throughput and automated pipelines have since improved success for tractable targets [14, 22].

### Prediction Tools and The Integrated Design Problem

Computational tools increasingly inform parts of this pipeline — such as structure prediction [6], stability-change (ΔΔ*G*) predictors, and post-translational-modification (PTM), functional-annotation, and disorder predictors [9, 10]. Yet benchmarks show these tools remain unreliable for the hardest cases [8], and, more importantly, the decision a scientist actually faces is *integrated* : a construct boundary interacts with tag placement, which interacts with expression host, which in turn interacts with purification behaviour. Even more importantly, these choices jointly shape the final *developability* — the solution-behaviour properties that decide whether purified protein is usable downstream (attainable concentration, monodispersity, and resistance to aggregation on handling [17]) — which can be distinct from raw expression yield. A prediction that improves one axis in isolation does not necessarily improve the molecule a team can carry forward (Figure 1). These choices are also constrained in practice: a given lab has only certain vectors, tags, hosts, and equipment on hand, so a viable design must be executable with what is available, not merely optimal in principle.

**Figure 1:**
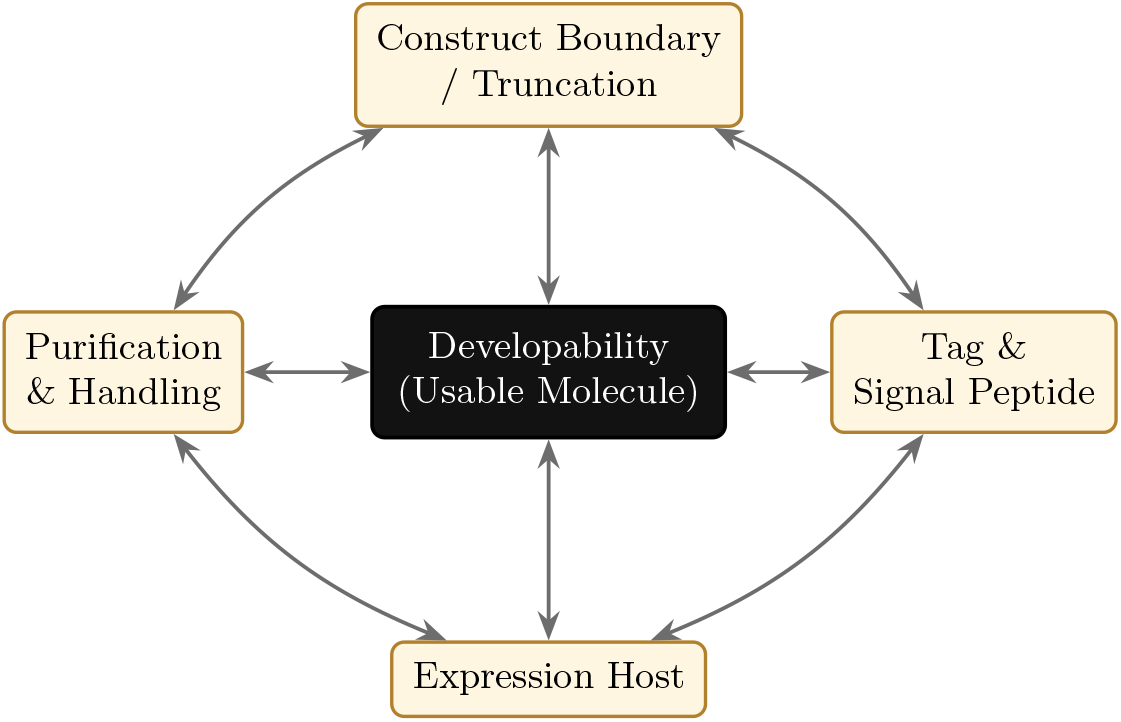
Interactions among construct-design decisions. Construct boundary, tag/signal peptide, expression host, and purification each interact with one another (outer links) and jointly determine developability (centre). Because the axes are coupled, a prediction that improves one in isolation need not improve the molecule a team can carry forward — which is why the platform scores and produces them together rather than one at a time.

Construct design itself has a smaller dedicated literature. Domain-boundary and construct-library methods (e.g. ESPRIT and related expressibility screens [13]) and disorder/secondary-structure-guided truncation are established for choosing where to cut, but they typically optimise expression or solubility in isolation rather than the joint outcome of expression *and* downstream developability, and they are not coupled to host-specific expression and purification protocols.

### Model Target: FGF21

We chose human fibroblast growth factor 21 (FGF21) as a focal, real-world target (Figure 2). FGF21 is a secreted endocrine hormone that regulates glucose and lipid metabolism and energy balance through the receptor tyrosine kinase FGFR1c and its obligate co-receptor *β*-Klotho [1, 2]. It is an active drug class: FGF21 analogues are in late-stage clinical trials for metabolic dysfunction-associated steatohepatitis (MASH), type-2 diabetes, obesity, and severe hypertriglyceridemia [3, 4]. It is also a challenging target for recombinant production and biophysical characterisation — a non-canonical *β*-trefoil core flanked by a long, disordered, proteolysis-prone C-terminal tail, with low intrinsic thermostability (reported melting temperature ∼46.8 °C [19]). The C-terminal tail is also functionally important: its distal end carries the *β*-Klotho binding determinants and is cleaved *in vivo* by fibroblast activation protein after Pro-199, which inactivates the hormone [21]. It is exactly the kind of stability-challenged target where construct design dominates outcome: recombinant FGF21 is commonly produced in *E. coli* with solubility-enhancing fusion partners (e.g. maltose-binding protein) or refolding [18, 19]; as a secreted endocrine factor it can instead be produced in a mammalian secretory host (the route taken here), underscoring how much the construct and process decide the result.

**Figure 2:**
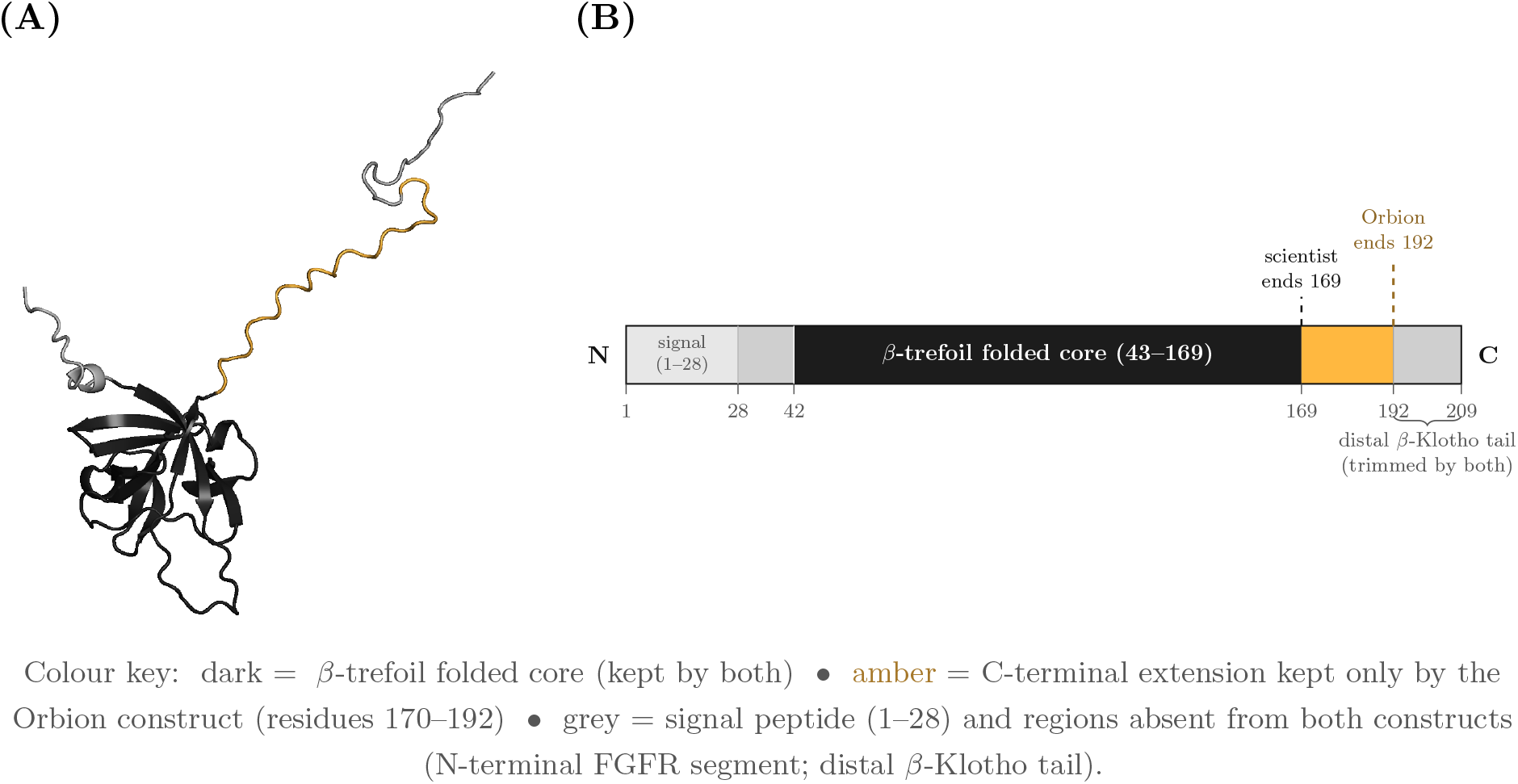
FGF21 Structure and the Two Construct Boundaries. **(A)** AlphaFold-predicted model of full-length human FGF21 (UniProt Q9NSA1; AlphaFold DB v6, 6, 20), coloured as the design map (UniProt Q9NSA1 numbering). The non-canonical *β*-trefoil core (dark; ≈43–169) is retained by both constructs and matches the experimental NMR structure of the core domain (PDB 6M6E, 5). The C-terminal tail is intrinsically disordered (low model confidence, a regime in which AlphaFold pLDDT is itself a disorder signal rather than a reliable conformation [7]); its precise modelled conformation is not meaningful and is shown only to convey the extent and disordered character of the retained region. **(B)**The same chain drawn as a to-scale domain architecture (segment widths proportional to residue counts), marking the FGFR1c-engaging N-terminus, the disordered *β*-Klotho tail, and the two construct end-points; at true scale the C-terminal tail is small relative to the folded core. The scientist construct ends at the folded-core boundary (residue 169); the Orbion construct extends ∼23 native residues further (to residue 192), retaining the proximal start of the tail (amber). Both constructs still remove the distal tail carrying the *β*-Klotho binding site and the N-terminal FGFR-engaging segment, so neither is intended to be receptor-active.

Residue numbering throughout refers to the full-length human FGF21 precursor (UniProt Q9NSA1, 209 aa): the signal peptide is residues 1–28 and the mature secreted chain is 29–209.

### Study Design

We ran a prospective, head-to-head wet-lab case study (Figure 3A) addressing two questions: (i) *did the platform’s prospective predictions match the experimental outcome*; and (ii) *did integrated, prediction-guided construct and protocol design yield a more usable molecule than the standard literature approach?* Two FGF21 expression constructs were produced side by side under the same experimental workflow, using two different design strategies. One was designed by a scientist from the literature, applying standard construct-design principles — the published folded-core boundary, a secretion signal, and an affinity tag — to balance expressibility, solubility, and downstream use; the other by the Orbion platform, an AI, prediction-guided system that additionally generated the expression and purification protocols. These are *workflow-level designs rather than single-variable constructs* — they differ in truncation boundary, signal peptide, and tag architecture — and we treat them as such throughout.

**Figure 3:**
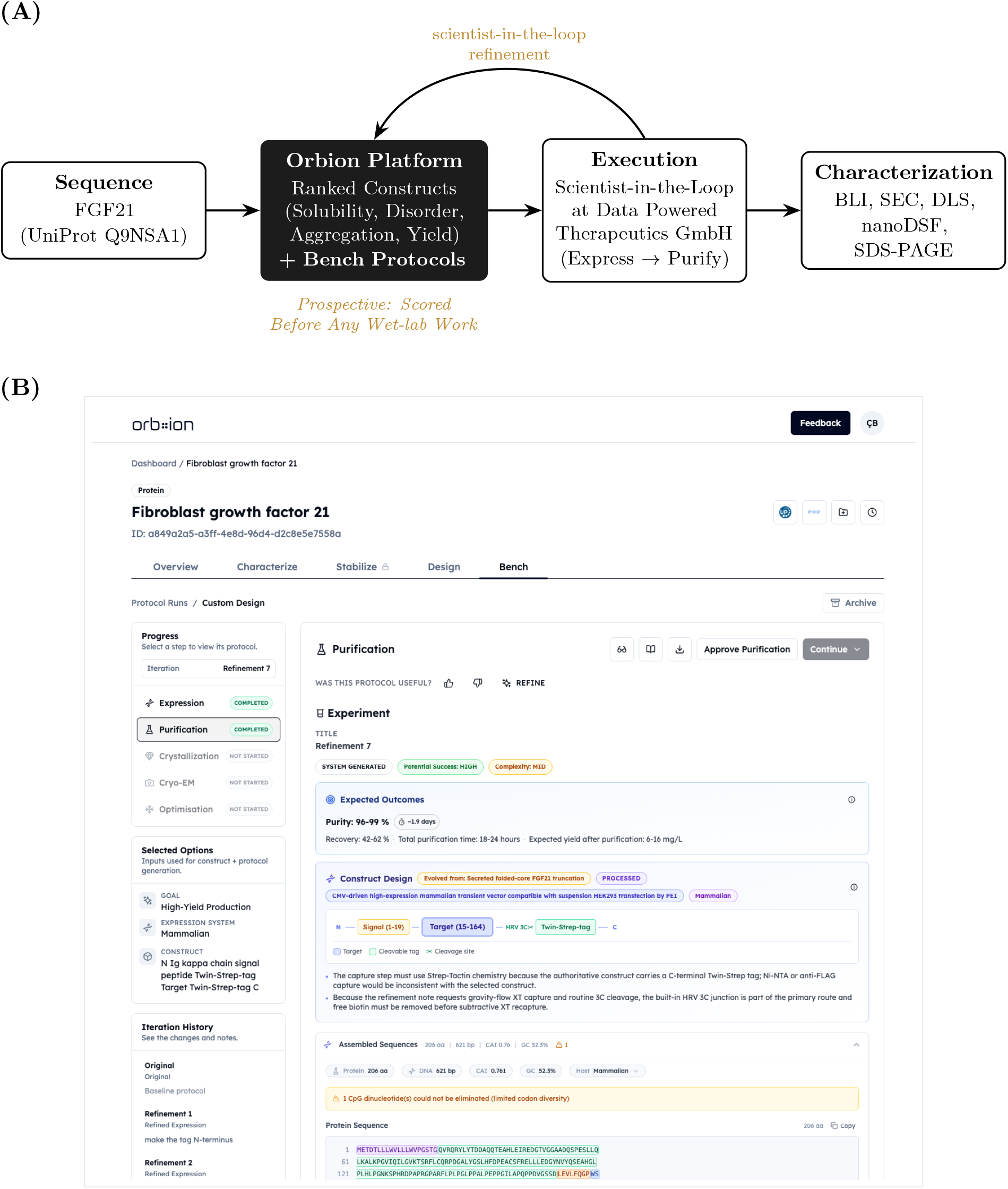
Design-to-bench workflow and a platform-generated protocol. **(A)** From a single sequence, the platform produced ranked construct designs *and* the expression/purification protocols. Both were executed scientist-in-the-loop at an external partner (Data Powered Therapeutics): the expression protocol was used essentially as generated; the purification followed the platform’s plan, refined at the bench. All platform scores were computed before the experiment. **(B)** A platform-generated Bench protocol (purification, “Refinement 7”) for the Orbion construct: the module emits human-readable, parameterised protocols with expected outcomes (here purity 96–99%, recovery 42–62%, expected yield 6–16 mg/L), the construct architecture (Ig-*κ* signal FGF21 target 15–164, i.e. UniProt 43–192· HRV-3C· C-terminal Twin-Strep), the assembled coding sequence, and an iteration history (left panel) recording the scientist-in-the-loop refinements.

## 2. Platform Design and Prospective Predictions

The Orbion platform uses protein sequence as input and returns ranked construct designs together with experimental conditions. The platform does not rely on a single end-to-end prediction alone. Instead, it assembles context from multiple specialist predictors — including solubility, disorder, aggregation, PTM liability, binding-region preservation, host-specific expression, and purification constraints — and uses these signals to rank construct designs and generate bench-executable protocols. The output is therefore a *developability-aware recommendation* rather than a single-property optimization.

We treat it here as a decision tool and report only the outputs used in this study; the underlying models — including AstraPTM2 for post-translational modifications [9], Astra-ROLE2/AstraSUIT2 for functional annotation and suitability [10], AstraBIND for ligand-binding sites [11], AstraUNFOLD for residue-level structural and disorder predictions, and AstraDTM for thermostability — are described separately and accessible at orbion.life/research. The expression and purification protocols it emits are *interactive, iterable* documents: a scientist refines them against bench reality and the platform regenerates accordingly (see Table 2).

### Construct predictions

For FGF21 the platform enumerated candidate constructs (signal peptide, tag architecture, fusion, and target truncation boundaries) and scored each construct with a set of per-property predictors — predicted solubility, intrinsic disorder, aggregation propensity, PTM liabilities, predicted binding sites, and expression yield — combined into a single ranking score (“composite”). Computed for the mammalian host used here, the composite scored the platform construct (termed “Orbion construct”) at **68.7** and the scientist construct at **59.0**.

### Construct rationale (verbatim platform output)

The platform was supplied with the *mature* FGF21 sequence (UniProt residues 29–209, which it numbers 1–181), so residue positions in the rationale below are in that mature frame: the selected boundary (mature 15–164) corresponds to UniProt 43–192. Throughout the paper the UniProt number is used, except for the direct output verbatim from the platform. The platform’s design rationale for the selected construct is reproduced below.

*Produce a prediction-guided core-domain construct that removes the long low-confidence/PTM-rich C-terminal tail beyond the high-confidence folded region. The structural analysis places a high-confidence domain at 15–142 and a core region at 15–164, so starting at 15 and ending at 164 removes >20 residues from each side of the most problematic extensions while retaining the folded extracellular core expected to drive binding-related function. Compared with literature-like full-length constructs, this should reduce secretion heterogeneity and proteolysis risk in mammalian medium, with the main trade-off being possible loss of any tail-dependent regulatory activity*.

### Bench protocols

The platform also generated the *expression* and *purification* protocols for the selected construct (Figure 3B): a transient mammalian (Expi293F/PEI) expression protocol with predicted yield and process parameters, and a Strep-Tactin XT capture *→* HRV-3C cleavage *→* size-exclusion polishing purification protocol. These were executed scientist-in-the-loop and iteratively refined (Section 4, Table 2).

## 3. The Two Construct Designs

Both constructs share an identical FGF21 core (UniProt Q9NSA1); they are compared as *workflow-level designs* and differ in truncation boundary, signal peptide, and tag architecture (Figure 4A, Table 1; see Limitations). The scientist construct reproduces the canonical folded-core boundary deposited as PDB 6M6E [5]; the Orbion construct retains ∼23 additional native C-terminal residues, a flexible, proline/serine-rich segment that is conventionally removed for expression and crystallisation (Figure 4B). A BLAST search against public sequence databases returned no construct with this boundary and tag architecture; the FGF21 sequence itself is 100% native (no engineered mutations), so the novelty is the design *choice*, not the sequence. Full amino-acid sequences used for gene synthesis are given in Appendix A.

**Table 1.** Construct design comparison.

|  | Scientist construct | Orbion AI construct |
| --- | --- | --- |
| Design Source | Literature / Expert Reading | Orbion Platform |
| FGF21 Core | Identical (UniProt Q9NSA1) | identical (UniProt Q9NSA1) |
| Truncation (UniProt) | 42–169 (Folded Core; = PDB 6M6E) | 43–192 (Core + ~23 Native Residues) |
| Signal Peptide | BM40 (SPARC) | Ig-κ |
| Tag Architecture | N-terminal Twin-Strep-3C | C-terminal 3C-Twin-Strep |
| Composite Score (Mammalian) | 59.0 | **68.7** |

**Table 2.** Scientist-in-the-loop refinement rounds on the platform-generated protocols.

| # | Stage | Bench feedback incorporated |
| --- | --- | --- |
| 1 | Expression | Move the affinity tag to the N-terminus |
| 2,4 | Expression | Replace ExpiFectamine with linear PEI as the transfection reagent |
| 3 | Expression | Use MEXi-293E cells (IBA) as the host line |
| 5 | Expression | Hold a constant 37 °C expression temperature |
| 6 | Purification | Add an HRV-3C tag-cleavage step after affinity capture |
| 7 | Purification | Use a gravity-flow column; biotin (not desthiobiotin) for Strep-Tactin XT elution; define HRV-3C cleavage buffer (20 mM Tris pH 7.5, 150 mM NaCl, 20 mM imidazole), target 0.5–1.0 mg/mL, and an explicit diafiltration buffer-exchange protocol |

**Figure 4:**
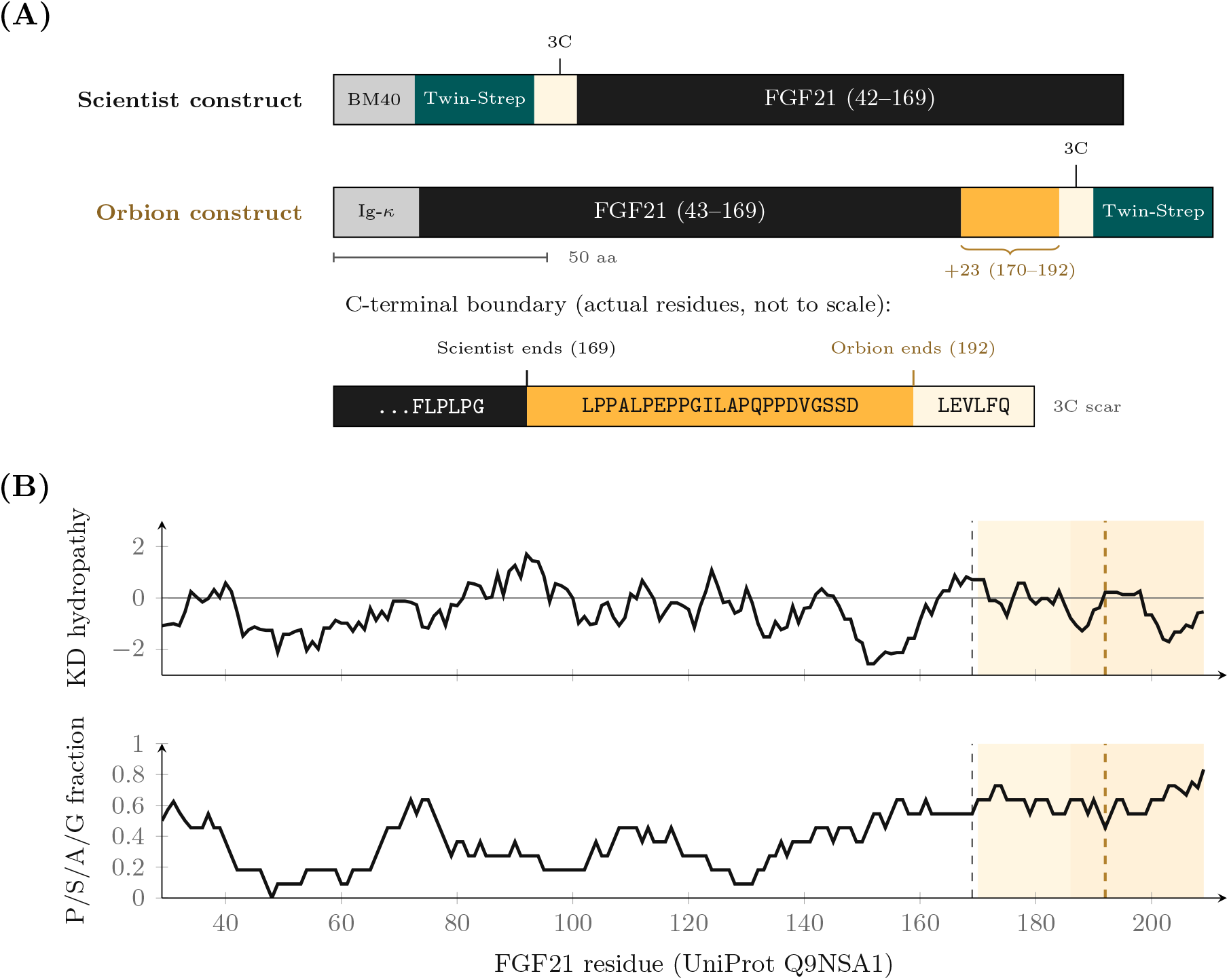
The two construct designs and their sequence context. **(A)** Construct architecture (N-terminus at left). Both share an identical FGF21 core (dark). The scientist construct truncates at the folded-core boundary (residue 169) and carries the Twin-Strep tag at the *N* -terminus; the Orbion construct retains a 23-residue native C-terminal extension (amber, residues 170–192) and carries the tag at the *C* -terminus. Signal peptides (grey; BM40 vs Ig-*κ*) are removed on secretion and the Twin-Strep tag (teal) is removed by HRV-3C cleavage (cream), leaving a short `LEVLFQ` scar at the C-terminus of the Orbion construct and a two-residue `GP` scar at the N-terminus of the scientist construct; the bottom strip shows the actual C-terminal boundary residues (not to scale), with the scientist (169) and Orbion (192) end-points marked. Bars are drawn to scale (shared 50-aa ruler). **(B)** Sequence-derived properties: top, Kyte–Doolittle hydropathy [23] (9-residue window); bottom, low-complexity content (fraction of Pro/Ser/Ala/Gly, 11-residue window). The disordered C-terminal tail (cream, residues 170–209) is strongly hydrophilic and low-complexity and carries the *β*-Klotho region (amber, ≈ 186–209); the scientist boundary (black dashed, 169) stops at the folded core, while the Orbion boundary (amber dashed, 192) retains the hydrophilic, Pro/Ser/Ala-rich segment 170–192. Residue numbering is UniProt Q9NSA1 throughout.

## 4. Methods

We report the protocols, including the parameters that were generated, those that were altered scientist-in-the-loop, and the conditions finally run. The platform-generated protocols were refined over seven rounds of bench feedback (Table 2). These refinements did not alter the purification *strategy* ; they aligned the protocol with the lab’s reagents, equipment, and standard practice (e.g. biotin — the correct elution reagent for Strep-Tactin XT with a Twin-Strep tag — rather than desthiobiotin) or added steps required for downstream characterisation (e.g. tag cleavage) — the same adjustments a scientist routinely makes when executing a purification at the bench. All platform construct rankings, per-property scores, and protocols reported here were generated and recorded *before* the constructs were synthesised; the head-to-head comparison is therefore prospective.

### Constructs and Gene Synthesis

Two synthetic genes encoding human FGF21 (UniProt Q9NSA1) were synthesised (Twist Bioscience) and cloned into a CMV-driven mammalian transient-expression vector. Plasmids were sequence-verified; full protein sequences are in the Appendix A, and shortly presented here: *Scientist construct* : BM40 signal peptide, N-terminal Twin-Strep tag, HRV-3C site, FGF21 residues 42–169. *Orbion construct* : Ig-*κ* signal peptide, FGF21 residues 43–192, C-terminal HRV-3C site and Twin-Strep tag.

### Expression (Platform Protocol, Scientist-in-the-Loop)

Transient expression was performed in suspension HEK293-derived cells (MEXi-293E, IBA; substituted for Expi293F per refinement, Table 2) in serum-free medium at a constant 37 °C, 5% CO_2_, 125 rpm. Cultures (30 mL) were transfected at 2.5–3.0× 10^6^ cells/mL (viability ≥95%) with 30 µg plasmid DNA and 90 µg linear polyethyleneimine (PEI) (25 kDa), 1:3 (w/w); DNA and PEI were each diluted in 1.5 mL growth medium, combined, incubated for 15–20 min at room temperature, and added dropwise. Secreted product was monitored across days 1–3 by bio-layer interferometry (BLI) on Strep-Tactin XT probes and the supernatant was harvested at day 3.5 by two-step centrifugal clarification (at 500*×g* for 5 min at room temperature and at 4,000*×g* for 1 h at 4 °C).

### Purification (Platform Protocol, Scientist-in-the-Loop)

Clarified medium was loaded by gravity onto 300 µL Strep-Tactin XT high-capacity resin (10 mL gravity column) and washed with 20 column volumes of 20 mM Tris-HCl pH 7.5, 150 mM NaCl, 20 mM imidazole. The protein was eluted by on-column HRV-3C cleavage (His-tagged 3C protease, 2 h at room temperature), collected in two 750 µL fractions. The eluate was passed over 200 µL Ni-NTA to remove the His-tagged protease (a lab-specific step added scientist-in-the-loop, since the protease’s His-tag was not part of the platform’s inputs), concentrated (Amicon Ultra-0.5, 3 kDa MWCO) to ∼450 µL, and polished by size-exclusion chromatography (SEC; Superdex 75 Increase 10/300, Cytiva). Peak fractions were pooled and concentrated to the final volumes (∼60 µL Orbion construct, ∼100 µL scientist construct) in 20 mM HEPES pH 7.5, 150 mM NaCl — the buffer in which all biophysical characterisation was performed. The platform-generated protocol additionally specified a subtractive Strep-Tactin recapture and a Q anion-exchange polish; the team executed the streamlined on-column-cleavage route above for comparability across the two constructs.

### Biophysical Characterisation

Final samples were concentrated and, before biophysics, clarified (10 min, 15,000*×g*, 4 °C). Yield and monodispersity were assessed by SEC; polydispersity index (PDI) and apparent melting temperature (*T*_*m*_) by dynamic light scattering (DLS) and nanoDSF (Prometheus Panta, NanoTemper Technologies; 10 acquisitions); and purity by SDS-PAGE (1.5 µg/lane; NEB #P7719 broad-range ladder). Protein concentrations were determined spectrophotometrically at 280 nm using sequence-derived extinction coefficients. All wet-lab work was performed at Data Powered Therapeutics GmbH.

### Statistics

This study compares a single construct pair produced in one expression/purification run (*n* = 1 per construct). No inferential statistical testing was therefore performed, and none is appropriate at this sample size. Where a spread is reported (DLS hydrodynamic radius, mean ± SD), it reflects ten repeat acquisitions of a single sample and characterises instrument/measurement precision, not biological or experimental replication. All cross-construct differences are reported descriptively.

## 5. Results

### Expression and Yield

Both constructs produced clean, secreted, Strep-capturable material. The scientist construct showed a faster bio-layer interferometry (BLI) on-rate at every time point and a higher purified yield: **0.71 mg vs 0.30 mg** from a 30 mL culture (∼24 vs ∼10 mg/L). Size-exclusion chromatography agreed — both eluted as single, symmetric peaks with no void aggregate, and the scientist construct peak was ∼3*×* larger, consistent with more material (Figure 5B). We report this opposite-of-favourable result first: on raw output, the scientist design won. The on-rate gap was consistent across the time course (response at 10 s: day 1, 1.1 vs 0.8 nm; day 2, 2.7 vs 1.3 nm; day 3, 3.5 vs 2.0 nm; scientist vs Orbion construct), while both plateaued near 4.2–4.3 nm at saturation (Figure 5A).

**Figure 5:**
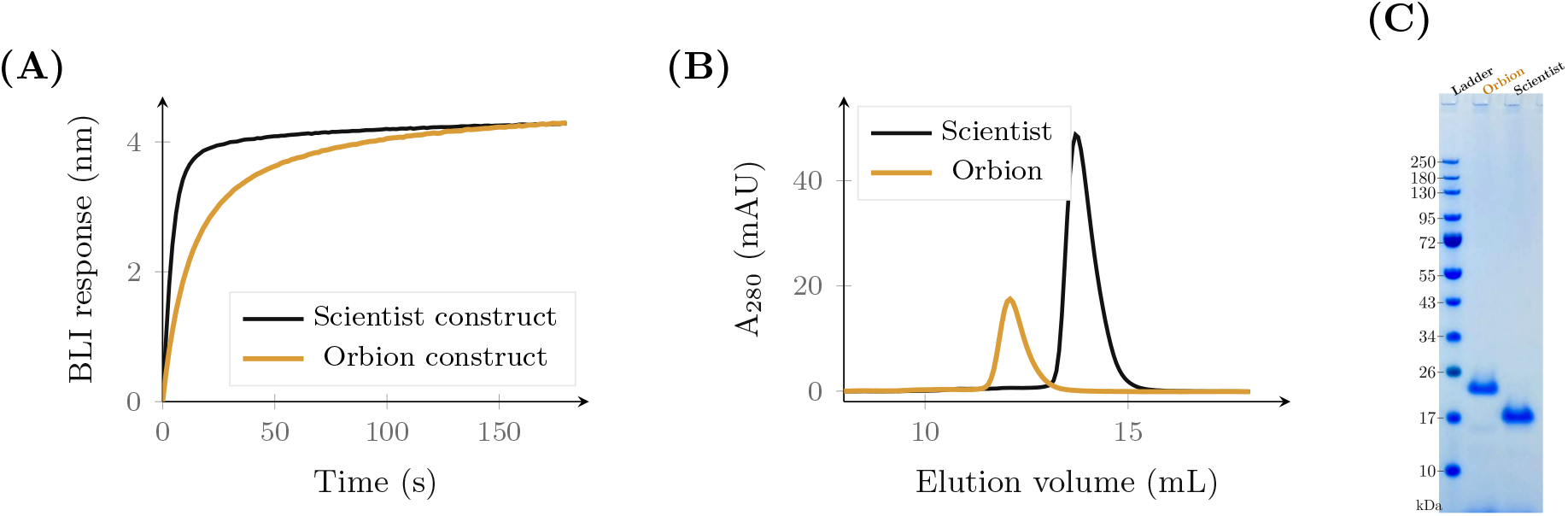
Expression, purification, and purity of the two constructs. **(A)** Expression time-course by bio-layer interferometry (day-3 supernatant): secreted, tagged protein binding Strep-Tactin XT probes. The scientist construct (black) shows a faster on-rate — a proxy for capturable secreted protein — consistent with its higher purified yield (response at 10 s is the informative readout; both plateau near saturation). **(B)** Size-exclusion chromatograms: both elute as single, symmetric peaks (no void aggregate); the scientist construct gives the taller peak (more protein), while the Orbion construct elutes earlier (12.1 vs 13.7 mL; larger apparent size), consistent with its longer native C-terminus. **(C)** Coomassie SDS-PAGE of the final samples (ladder; Orbion; Scientist; 1.5 µg/lane), run after SEC and concentration: both migrate as a single clean band, confirming high purity.

### Developability and Solution Behaviour

The constructs diverged at the concentration step. **The Orbion construct concentrated cleanly to 1.4 mg/mL**, while **the scientist construct stalled at 0.7 mg/mL**, losing material to aggregation and/or adsorption to the concentrator membrane as it was concentrated.

This means the scientist construct’s ∼2.4*× purified-yield* advantage did not survive concentration: although it entered the concentration step with substantially more protein (its SEC peak was ∼2.5*×* taller; Figure 5B), in the final concentrated samples the Orbion construct held *more* usable protein (∼84 µg, 1.4 mg/mL in ∼60 µL) than the scientist construct (∼70 µg, 0.7 mg/mL in ∼100 µL). The full SEC peak was concentrated for both (not an aliquot), so this reflects a loss of the scientist construct’s material on concentration. These are single, approximate end-point measurements; the robust differentiators are the attainable concentration and monodispersity, not the exact microgram amounts. We evaluated this monodisperse behaviour by DLS, in which the Orbion construct was more monodisperse (lower PDI, 0.21 vs 0.30, although neither sample is strictly monodisperse in the DLS sense; Figure 6B). A DLS thermal-ramp aggregation assay was also attempted, but its high-temperature signal was a transient spike consistent with a bubble/degassing artifact rather than protein aggregation (the acquisitions were also flagged for inconsistencies), so we do not interpret it and do not use it as a developability readout. Assessing their purity, both were pure single bands by SDS-PAGE (Figure 5C) — so the Orbion construct’s advantage is *not* “dirty but soluble.” As a further indicator of protein stability, melting temperatures were measured by Nano Differential Scanning Fluorimetry (nanoDSF; Prometheus Panta, NanoTemper Technologies). Apparent melting temperatures were indistinguishable (∼43.6–43.9 °C) and, given FGF21’s low tryptophan content, could not be assigned with confidence (Figure 6A). All measured quantities are summarised in Table 3 and Figure 7.

**Table 3.** Biophysical characterisation summary. DLS cumulant radius is reported as mean ± SD over 10 acquisitions.

| Property | Scientist construct | Orbion construct |
| --- | --- | --- |
| Purified Yield | 0.71 mg ($\sim 24$ mg/L) | 0.30 mg ($\sim 10$ mg/L) |
| SEC Retention Volume | 13.7 mL | 12.1 mL |
| SEC Peak Area (Relative $A_{280}$) | $\sim 3.0$ | 1.0 (ref.) |
| Hydrodynamic Radius $R_h$ (nm) | $1.78 \pm 0.26$ | $2.34 \pm 0.10$ |
| Polydispersity Index | 0.30 | 0.21 |
| SLS Intensity (cts/s) | $1.05 \times 10^6$ | $1.47 \times 10^6$ |
| Signal/Noise (DLS) | 388 | 3074 |
| Max. Attainable Concentration | 0.7 mg/mL | 1.4 mg/mL |
| Final Concentrated Amount<sup>‡</sup> | $\sim 70$ μg | $\sim 84$ μg |
| Apparent $T_m$ (nanoDSF)* | $\sim 43.6$ °C | $\sim 43.9$ °C |
| Purity (SDS-PAGE) | Single Dominant Band | Single Dominant Band |

Green marks the construct that clearly leads on each property — a descriptive direction from a single construct pair ( $n = 1$ , one run), *not* a statistical inference (no significance testing is possible at this $n$ ; the $R_h \pm$ SD reflects ten repeat acquisitions of one sample). Unshaded rows are ties, or differences that are structural consequences of the Orbion construct’s longer native C-terminus (larger apparent size, earlier elution, higher scattering) rather than quality differences.
\*Apparent transition only; the low tryptophan content precludes a confident $T_m$ assignment for either construct.
<sup>‡</sup>Protein recovered in the final concentrated sample (concentration × volume: $\sim 0.7$ mg/mL × $\sim 100$ μL vs $\sim 1.4$ mg/mL × $\sim 60$ μL); approximate, single end-point. Although the scientist construct gave $\sim 2.4\times$ more purified protein, this advantage did not survive concentration: it lost material to aggregation and ended below the Orbion construct in usable concentrated form.

**Figure 6:**
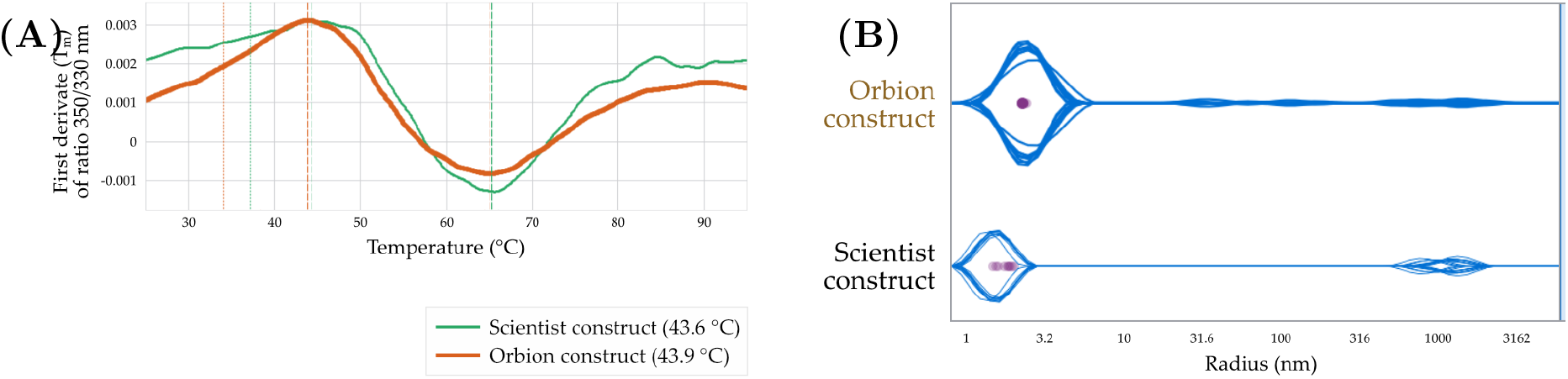
Thermal unfolding and solution monodispersity. **(A)** nanoDSF thermal unfolding (first derivative of the 350/330 nm ratio): an apparent transition near 43–44 °C is seen for both constructs (apparent *T*_*m*_ 43.6 vs 43.9 °C), but the low tryptophan content makes the signal shallow and the true *T*_*m*_ cannot be confidently assigned — consistent with FGF21’s reported low intrinsic thermostability. **(B)** DLS size-distribution analysis (Prometheus Panta): the Orbion construct is low-polydispersity across ten acquisitions (PDI 0.21, high signal/noise), whereas the scientist construct is more polydisperse (PDI 0.30) and shows a more prominent high-molecular-weight population near 1 µm — the heterogeneity underlying its concentration behaviour.

**Figure 7:**
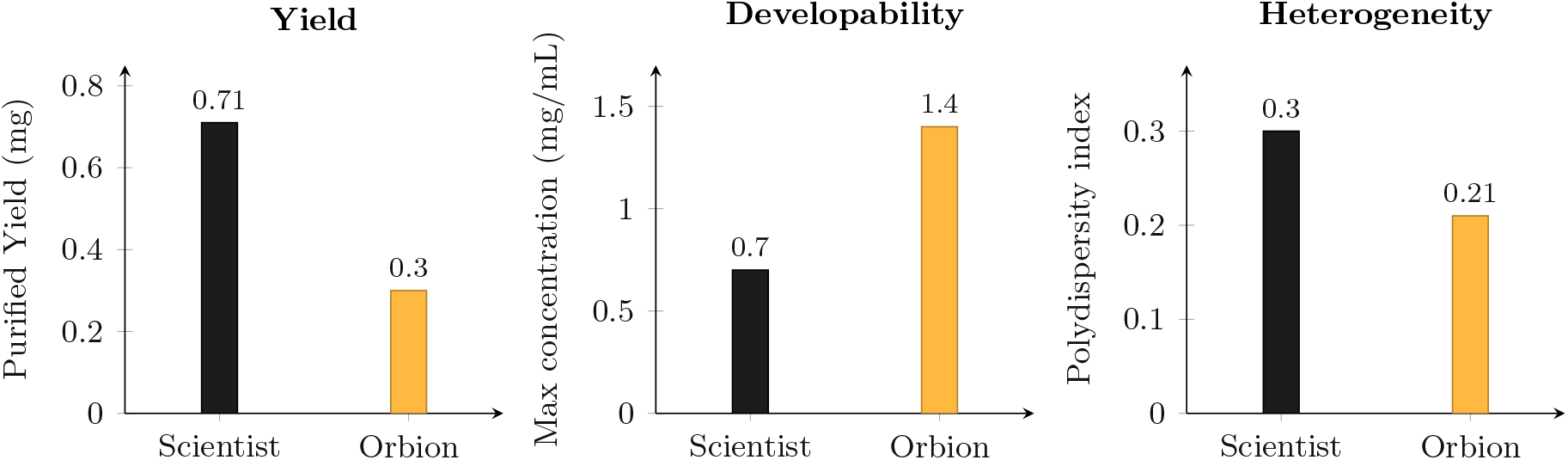
Yield and developability of the two constructs. The scientist construct gave more total *purified* protein (left) — but that advantage did not survive concentration (the Orbion construct held more usable protein in the final concentrated sample; Table 3). The Orbion construct reached twice the attainable concentration without aggregating (centre) and was more monodisperse (lower PDI; right) — the properties that determine whether a sample is usable downstream. Amber = Orbion AI design.

### Prospective Predictions versus Outcome

Computed for the mammalian host used, the platform’s composite score had ranked the Orbion construct above the scientist construct (**68.7 vs 59.0**) *before* the experiment. Its predictions of purified yield, solubility, and disorder also matched what the lab observed (Table 5). The underlying per-property predictor outputs are listed in Table 4: notably, the higher predicted intrinsic disorder for the Orbion construct (27.6% vs 13.5%) is consistent with its retained, disordered C-terminus (Figure 4B), and predicted solubility was high for both. The aggregation sub-score did not discriminate the two constructs (and, at face value, slightly favoured the scientist construct), and the thermostability prediction (Δ*T*_*m*_) could not be tested. We therefore make a bounded claim: the platform’s *ranking and several of its property predictions were correct in advance*; we do not claim the per-residue mechanism was proven (see Limitations).

**Table 4.** Platform per-property predictions for the two constructs. Per-property values are individual predictor outputs; the composite is the platform’s developability ranking score.

| Predictor (model) | Scientist construct | Orbion construct |
| --- | --- | --- |
| Solubility (%) | 99.0 | 99.3 |
| Intrinsic Disorder (%; AstraUNFOLD) | 13.5 | 27.6 |
| Aggregation Propensity (%; AstraUNFOLD) | 49.3 | 50.2 |
| **Composite (Mammalian Host)** | **59.0** | **68.7** |

**Table 5.** Prospective platform predictions vs. wet-lab outcome.

| Axis | Prediction (Mammalian Host) | Wet-lab Outcome |
| --- | --- | --- |
| Composite Ranking | Orbion > Scientist (68.7 vs 59.0) | Orbion More Developable |
| Purified Yield (Orbion) | 6–16 mg/L | ~10 mg/L |
| Solubility | Both Soluble (~99%) | Both Soluble (Non-discriminating) |
| Disorder | Orbion More Extended | Larger Apparent Size, Longer C-term |
| Aggregation | ~Equal (50.2 vs 49.3) | Scientist More Prone |
| Thermostability | $\Delta \sim 0.3^\circ\text{C}$ | Unassignable (low Trp) |

Green = the platform’s prospective prediction matched the wet-lab outcome. Solubility matched but was non-discriminating (both soluble). Aggregation was the one non-match — the sub-score was essentially equal and, at face value, slightly favoured the scientist construct, opposite to its greater aggregation on concentration; thermostability could not be assigned (low tryptophan).

#### Summary of results

The platform generated a novel, folded construct, as well as the according expression and purification protocols, executed scientist-in-the-loop. The platform had scored its own design higher than the scientist construct in advance (68.7 vs 59.0), and its yield/solubility/disorder predictions were confirmed. Under the same experimental workflow, the scientist construct expressed more protein, but the platform-designed Orbion construct showed a more *developable* profile — reaching twice the usable concentration and staying more monodisperse — ultimately being a more favourable construct candidate.

## 6. Discussion

This case study supports a specific, bounded claim: in this single-pair comparison, the Orbion platform-designed construct showed a more favourable downstream developability profile than the literature-derived construct, despite lower expression yield; the platform also generated the expression and purification protocols and scored its own design higher in advance. The platform did not simply reproduce the conservative, published construct; it proposed an *unconventional boundary* — keeping a flexible C-terminal segment that the field generally truncates — and that choice produced the better-behaved molecule. This is the behaviour that distinguishes a decision aid from a faster search: exploring design choices that human heuristics tend to exclude.

For the teams this work is aimed at, the practical reading is that the highest-leverage decision is often not the obvious one (maximise yield) but the developability-aware one (a molecule that survives concentration and handling). In this case the developability advantage was large enough to *reverse* the apparent yield gap in practical terms: although the scientist construct gave ∼2.4*×* more *purified* protein, it shed material on concentration, so in the final usable concentrated sample the Orbion construct delivered more protein (∼84 vs ∼70 µg; Table 3). Raw purified yield therefore overstated the scientist construct’s practical advantage — the usable amount, at usable concentration, favoured the prediction-guided design. (These are single, approximate end-points; the durable differentiators remain attainable concentration and monodispersity.) A platform that scores constructs on this axis, and that also produces executable, refinable protocols, addresses the bottleneck where programmes regularly stall.

### A biophysical hypothesis

The retained segment (residues 170–192) is hydrophilic and Pro/Ser/Ala-rich (Figure 4B) — the signature of an intrinsically disordered, low-complexity region. Such segments are known to behave as “entropic bristles” or built-in solubility enhancers: they enlarge the hydrodynamic radius and can shield aggregation-prone surfaces, the same role exploited by disordered fusion solubility tags. This is consistent with the properties we measured for the Orbion construct — a larger *R*_*h*_, earlier SEC elution, and cleaner concentration to higher levels — and rationalises why retaining a boundary the field usually trims *improved*, rather than harmed, developability.

We are cautious about mechanism, but the comparison is cleaner than it first appears. Because the affinity tag is cleaved before concentration (and the signal peptide is removed during secretion), the mature proteins measured at the concentration step differ principally in the retained C-terminal region — so the tag *position* is unlikely to explain the developability difference, as the wet-lab partner independently noted. That points to the retained C-terminal segment itself: a Pro/Ser-rich, disordered extension that can improve solubility, possibly together with the residual C-terminal protease scar. Two candidate effects — a solubilising disordered tail and reduced exposure of a proteolysis-prone region — remain hard to separate at *n* = 1, and we cannot fully exclude folding-history effects from the differing expression-stage architectures. The honest statement is that the design choice paid off and is most consistent with the retained C-terminus, not that a single causal hypothesis was proven.

## 7. Limitations

We state the limitations plainly, because they bound what may be concluded:

- **Single pair, single run (***n* = 1). This is a case study, not a statistically powered benchmark; the reported effect is a demonstration of capability, not an estimate of a population mean.
- **A single, partly-confounded pair**. The constructs differ in C-terminal boundary, signal peptide, and tag position. However, the signal peptide is removed during secretion and the affinity tag is removed by on-column 3C cleavage *before* the developability measurements, so the mature proteins compared at the concentration step differ principally in the retained C-terminal region. The affinity-tag *position* therefore cannot account for the concentration behaviour, and the developability advantage is most consistent with the retained C-terminal extension — though at *n* = 1 we cannot fully exclude folding-history effects from the differing expression-stage architectures or the small residual protease scars.
- **Buffer-specific behaviour**. The solution behaviour reported here was measured in a single formulation buffer; concentration and aggregation behaviour are buffer-dependent, and a different formulation could shift the comparison.
- **Biological activity not assayed**. “Developable” here means solution behaviour, not function; both constructs trim FGF21’s receptor-engaging regions.
- **Measurement caveats**. The DLS comparison was noisier for the scientist construct, and apparent *T*_*m*_ could not be confidently assigned for either construct.
- **Identity not confirmed by mass spectrometry**. Purity was assessed by SDS-PAGE; we did not acquire intact-mass (ESI-MS) data to confirm construct identity and integrity, which would be standard before downstream use.

## 8. Conclusion

For difficult proteins, the bottleneck is upstream of the biology — in construct design, expression, and purification, which are still decided largely by precedent and intuition and which fail for most hard targets. In a prospective, head-to-head wet-lab case study on FGF21, the first synthesised Orbion platform construct expressed and yielded usable protein with no make-test-redesign cycle, and furthermore showed a more favourable downstream developability profile than the literature-derived construct, despite lower raw yield. The platform also generated the protocols to produce it and had scored its own design higher in advance — choosing an unconventional boundary the field generally avoids. The result is a single, scoped data point, but it demonstrates a capability relevant to a costly bottleneck. Prediction-guided, developability-aware construct and protocol design is a plausible lever to remove iteration cycles and to make otherwise-stalled targets tractable; quantifying that effect across many targets is the natural next step.

## Acknowledgments

The wet-lab study was carried out at **Data Powered Therapeutics GmbH**. We thank **Aparna Pottikkadavath**, who used the Orbion platform to generate and evaluate the construct designs and who created and iteratively refined the expression and purification protocols at the bench, and **Dr. Nikolay Dobrev**, who oversaw the full experimental process — both for their careful work and for an honest, independent reading of the results.

## Author Contributions

Çağlar Bozkurt and Evangelia Nathanail conceived and designed the experiment, conducted the follow-up discussions with the team at Data Powered Therapeutics GmbH, led the analysis and interpretation, and wrote the manuscript. Aniruddh Goteti supported the work by observing the experimental process and cross-checking specific technical details. All authors reviewed and approved the final manuscript. Generation and evaluation of the construct designs, development and iterative refinement of the protocols, and execution of the wet-lab experiments were carried out at Data Powered Therapeutics GmbH, as detailed in the Acknowledgments.

## Data and Code Availability

Raw instrument data (BLI, SEC, DLS/nanoDSF, SDS-PAGE), construct sequences, and the platform-generated protocols are available on request. The platform is accessible at orbion.life.

## A. Construct Sequences

**Orbion AI construct** (Ig-*κ* signal *·* FGF21 43–192 *·* HRV-3C *·* Twin-Strep; 206 aa):

~~~
METDTLLLWVLLLWVPGSTGQVRQRYLYTDDAQQTEAHLEIREDGTVGGAADQSPESLLQ
LKALKPGVIQILGVKTSRFLCQRPDGALYGSLHFDPEACSFRELLLEDGYNVYQSEAHGL
PLHLPGNKSPHRDPAPRGPARFLPLPGLPPALPEPPGILAPQPPDVGSSDLEVLFQGPWS
HPQFEKGGGSGGGSGGSAWSHPQFEK
~~~

**Scientist construct** (BM40 signal *·* Twin-Strep *·* HRV-3C *·* FGF21 42–169; 185 aa):

~~~
MRAWIFFLLCLAGRALASAWSHPQFEKGGGSGGGSGGSAWSHPQFEKSGLEVLFQGPGQV
RQRYLYTDDAQQTEAHLEIREDGTVGGAADQSPESLLQLKALKPGVIQILGVKTSRFLCQ
RPDGALYGSLHFDPEACSFRELLLEDGYNVYQSEAHGLPLHLPGNKSPHRDPAPRGPARF
LPLPG
~~~

